# Alteration of Substrate-Induced Conformational Changes of *Escherichia coli* Melibiose Permease by Mutating Arg149

**DOI:** 10.1101/2020.05.31.125815

**Authors:** Yibin Lin

## Abstract

Fourier transform infrared difference spectroscopy and fluorescence spectroscopic techniques have been used to obtain information about substrate-induced structural changes of the melibiose permease mutant R149C, compared with the Cys-less, which were reconstituted into liposomes. ATR-FTIR evidences show that Na^+^-induced difference spectra of R149C and Cys-less are similar. However, Na^+^ induces some new peaks for R149C mutant permease. This means that replacement of Arg-149 by Cys may affect the structure of MelB, and then affect the binding of Na^+^. Melibiose-induced difference spectra of R149C in the presence of Na^+^ show some peaks in the amide I region not seen in Cys-less, corresponding to turns, β-sheets, α-helix changes. This suggests that R149C mutant permease undergo some different secondary structure changes compared to Cys-less mutant permease, when binding melibiose. Comparison of the permease intrinsic fluorescence variations of R149C and Cys-less indicate that there are similar substrate binding properties between R149C and Cys-less. When analyzing the effects of different sugars it appears that the R149C mutant is more sensitive to the sugar. All these data indicate that replacement of Arg-149 by Cys will affect Na^+^ and sugar binding, and enhance the selectivity and sensitivity to sugars.

## Introduction

Melibiose permease of Escherichia coli (MelB), encoded by the *melB* gene in the mel operon ^1^, is a well-studied representative of the glycoside-pentoside-hexuronide /cation symporter family of membrane transporters ^2^. It consists of 473 amino acids (molecular mass, 53 kDa), 70% of which are apolar ^3,4^. MelB utilizes free energy released from the energetically downhill movement of a cation (Na^+^, Li^+^, or H^+^), in response to an electrochemical cation gradient, to drive the uphill stoichiometric accumulation of a galactopyranoside ^5–7^. MelB binds and transports melibiose and its coupling ion in a 1:1 ratio ^7^. The three coupling ions have a common binding site, and Na^+^ or Li^+^ ions enhance the cotransporter affinity for melibiose ^8–11^.

Biochemical studies, including immunological, PhoA-MelB fusions, and proteolytic mapping analysis ^12–14^, as well as two-dimensional crystallization ^15,16^ and Fourier transform infrared (FTIR) studies ^17^, consistently fit a topological model that includes 12 transmembrane α-helical domains and with the N and C-terminal located in the cytoplasm. This secondary structure model is illustrated in Fig. 1. Moreover, genetic and site-directed mutagenesis studies ^3,4,18–20^ suggest that several acidic residues located in membrane domains of the N-terminal half of MelB may form a coordination network involved in ion recognition and the sugar-binding site may be preferentially located in the C-terminal half of MelB, and the N and C domains may be close to each other in 3D structure. It has also been suggest that helix IV may connect the two substrate-binding sites ^20^, and loops 4-5 and 10-11 are important for MelB function ^13,21–23^; and 3), Asp and Glu residues are implicated in cation and melibiose binding and/or translocation ^4,22,24–26^.

**Figure 1.**
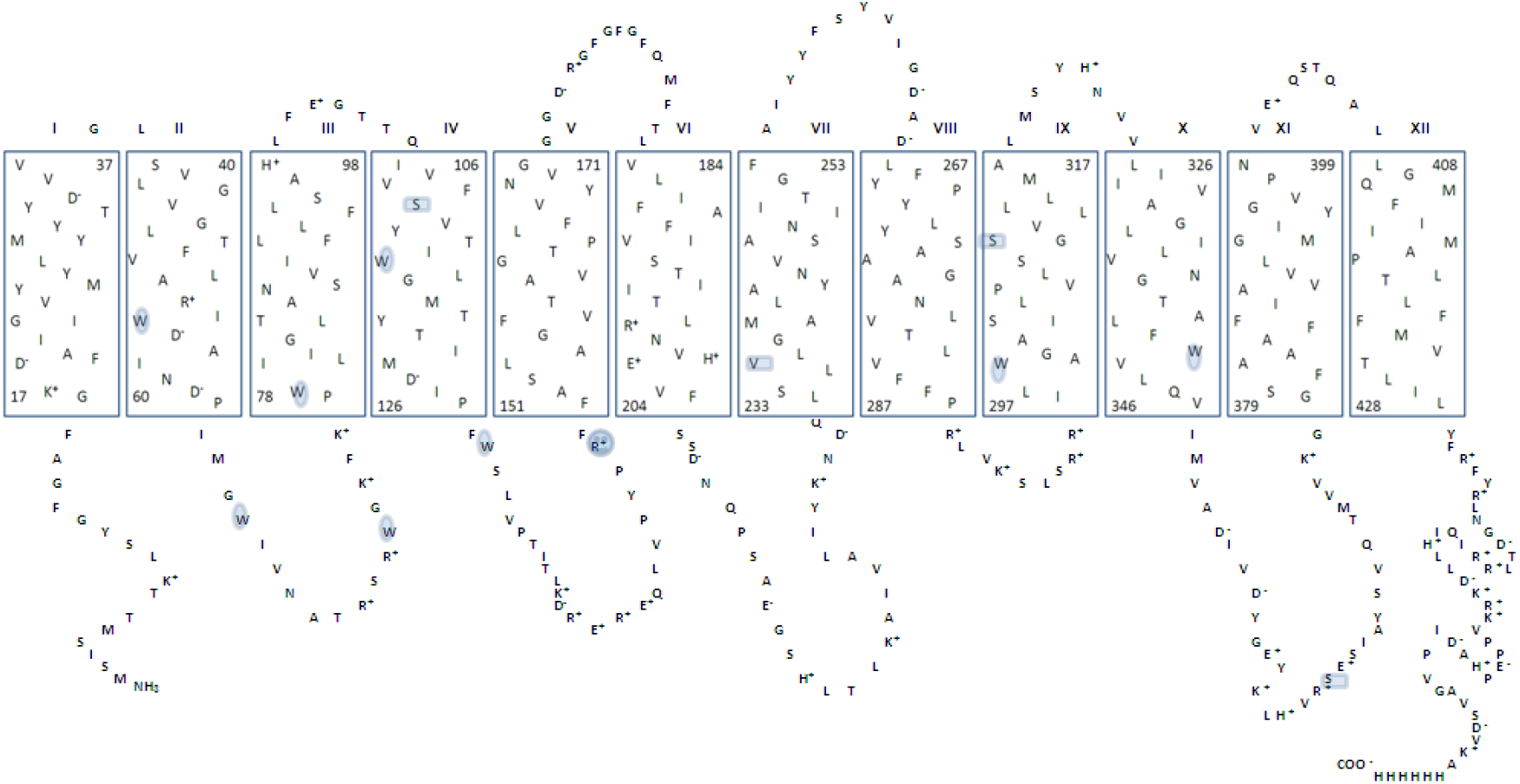
Predicted secondary structure of MelB. “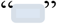” show the positions of four native cysteines which were mutated into Ser and Val in Cys-less mutant. Amino acid mutated into cysteine in this work is marked using “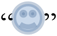”. All tryptophans are marked with “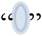”. The picture is taken from Pourcher *et al.* [Pourcher *et al.* 1996]

One of the most important aims of protein research is to study the relation between structure and function. While awaiting improved resolution of the MelB crystal structure, other techniques are being used to understand better the functioning of this transporter and identify amino acids involved in substrate binding and/or translocation. Cysteine-scanning mutagenesis is now a widely used strategy to study systematically the structure and structure-function relationship of membrane proteins. It is currently applied to MelB using a fully functional active permease devoid of its four native cysteines (Cys-less mutant permease) ^27^. as genetic background. Recently attenuated total reflection (ATR)-FTIR difference spectroscopy has been applied to study the substrate-induced changes in MelB conformation during the MelB transport cycle ^28–31^. Results from FTIR spectroscopy have not only provided information on the secondary structure components of MelB (Dave et al. 2000), they also suggest that changes in α-helix tilting occur upon substrate binding ^32^. An electrophysiological approach revealed fast transient currents (20-ms range) triggered by either Na+ or melibiose binding, resulting from movements of charged amino acids and/or a reorientation of helix dipoles ^33^. Intrinsic fluorescence spectroscopy has been used to investigate the effects of sugar and coupling ions on the conformation of MelB permease ^34^. While Na^+^ and Li^+^, on their own, quench MelB permease fluorescence, sugars enhance it, and the intensity of this increase is significantly potentiated by addition of the monovalent ions. These and other results led to the suggestion that fluorescence signal increase monitors changes of the transporter conformation associated with binding of the substrates.

Former studies have showed that loop IV-V is either directly involved in Na+ and /or sugar binding or contributes indirectly to the coupling interaction between the two binding sites. Cys-less replacement analysis indicated that Arg-141 and Arg-149 are essential for MelB transport activity. Replacing Arg-149 by a neutral Cys residue leads to selective inactivation of MelB translocation capacity ^21^. Biochemical analyses have shown that the R149C mutant was completely inactive ^21^. As well as a fluorescence resonance energy transfer signal arising from a bound dansylated sugar-analog has shown that this mutant does not retain Na^+^-dependent sugar-binding activity ^21^.

To understand the role of Arg-149 in the MelB, we sought to determine the extent to which the R149C defect is associated with any structural defect. To that end, we carried out a difference ATR-FTIR analysis and intrinsic fluorescence variations of the cosubstrate-induced change recorded on the purified R149C MelB mutant in proteoliposomes. The overall difference ATR-FTIR and intrinsic fluorescence variations informations obtained from MelB Cys-less mutant (the quadruple mutant C110S/C235V/C310S/C364S, 40) were used as a general reference. Overall, the data show that the R149C mutant introduces a selective defect of the structural properties of MelB during binding of substrates.

## Materials and Methods

### Materials

3-(laurylamido)-*N,N*’-dimethylaminopropylamine oxide (LAPAO) and n-dodecyl beta-D - maltoside (DDM) were obtained from Anatrace. Ni-NTA resin was obtained from SIGMA^®^. SM-2 Bio-Beads was from Bio-Rad Lab., Inc. Total E. coli lipids (acetone/ether precipitated) were purchased from Avanti Polar Lipids, Inc. AEBSF (4-(2-Aminoethyl) benzenesulfonyl fluoride hydrochloride) was from Melford Laboratories Ltd. High purity grade salts or chemicals (Suprapur, Merck) were used to prepare nominally Na^+^-free media containing less than 20 μM sodium salts. All other chemicals were obtained from commercial sources.

### Bacterial strains and plasmids

*E.coli* DW2-R, a *recA^−^* derivative of strain DW2 (*mel*A^+^, Δ*mel*B, Δ*lac*ZY) was used throughout the whole study ^35^. This strain is devoid of internal melibiose and lactose permease, but encodes for α-galactosidase. Preparation of competent cells was achieved by treating *E.coli* with calcium and rubidium chloride.

The vector pK95ΔAHB, which carries an ampicillin resistance and encodes a permease devoid of its four native cysteines (Cys-less permease) was used as matrix. This Cys-less carrier contains a valine instead of Cys-235 and serines instead of Cys-110, Cys-310, and Cys-364 ^27^. The vector of R149C was reconstituted based on this Cys-less carrier replacing Arg-149 by neutral Cys residue ^36^. These vectors which carry Cys-less mutant and R149C mutant were friendly given by Prof. Leblanc. *melB* is terminated by six successive triplets encoding His residues at its 3’ extreme. All mutants were generated as described in ^37–39^.

### Bacterial culture

*E.coli* DW2-R bacteria were transformed with the plasmid containing Cys-less or R149C mutated *melB* by using a heat-shock at 42°C ^40^. Transformed cells were plated on MacConkey agar plates (containing 10 mM melibiose and 100 μg/mL ampicillin) and incubated overnight at 30°C.

When melibiose was transported by the permease, it was hydrolyzed by α-galactosidase (encoded by the *melA* gene) into glucose and galactose. In response to the acidic by-products of sugar metabolism, the respective colonies turned red. In absence of melibiose transport, e.g., if a mutated transport-deficient *melB* gene was transformed into the bacteria, white colonies were observed.

Cells were selected and incubated at 30°C in Luria-Broth (LB) rich medium (12 mL, 10 g/L bacto tryptone, 5 g/L bacto yeast extract, 5 g/L NaCl). 100 μg/mL ampicillin and 10 μg/mL tetraciclin were added to the medium in order to maintain the selection pressure. This pre-culture was used either to isolate plasmid DNA for sequencing or as inoculate for large scale cultures.

In the latter case, cells (Cys-less or R149C mutant permease) were incubated in 200 mL M9 minimal medium (6 g/L Na_2_HPO_4_, 3 g/L KH_2_PO_4_, 0.5 g/L NaCl, 1 g/L NH_4_Cl) with the addition of 5 g/L glycerol, 1 mM MgSO_4_, 0.1 mM CaCl_2_, 1 mM thiamine, 2 g/L casamino acid, and 100 μg/mL ampicillin, 10 μg/mL tetraciclin. Cells were grown at 30°C under continuous shaking until an OD_600_ of 1-1.2 was reached. Large scale cultures (4 L) were carried out at big incubator. After overnight growing, cells were harvested, resuspended in 50 mM Tris buffer, pH 8.0 containing 50 mM NaCl, 5 mM β-mercaptoethanol, and directly used, or stored in the presence of 15% glycerol at −80°C until further use.

### Protein purification

Inverted membrane vesicles (IMVs) were prepared using a Microfluidizer Processors [M-110S, Microfluidizer^®^]. All steps were carried out at 0-4°C. Approximately 60 g (wet weight) of Cys-less and R149C mutated bacteria were rapidly thawed at 30°C and rinsed several times in a buffer containing 50 mM Tris-HCl (pH 8.0), 50 mM NaCl, and 5 mM β-mercapto-ethanol (resuspension buffer). Cells were resuspended in 100 mL of the same buffer supplemented with 5 mM EDTA for destabilization of the external bacterial membranes. Directly before applying a pressure of 20000 psi, 20 μg/mL of each DNAse and RNAse as well as 15 mM MgSO4 were added to the cell suspensions. An excess of EDTA (15 mM) was then added to the product which passed microfluidizer and unbroken cells and large debris were eliminated by centrifugation at 4000 × g (Beckman Coulter™, USA). The supernatant containing the IMVs was collected and washed several times with resuspension buffer by centrifugation at approximately 310,000 × g at 4°C for 30 minutes (SORVALL^®^ Centrifuges, USA). Finally, vesicles were resuspended in the same buffer (final volume no more 25 mL), frozen the sample using liquid N2 and stored at −80°C until further use.

Protein purification was carried out mainly as described ^41^. IMVs were thawed in water bath at 30°C and washed once in 600 mM NaCl, 20 mM Tris-HCl (pH 8.0), 5 mM β-mercaptoethanol, and 10 % glycerol (buffer I). All following steps were performed at 0-4°C. After centrifugation at 310,000 × g for 30 minutes (SORVALL^®^ Centrifuges, USA), the pellet was resuspended in a buffer of 10% glycerol, 600 mM NaCl, 20 mM Tris-HCl (pH 8.0), 5 mM β-mercaptoethanol, 10 mM imidazole, and 10 mM melibiose (buffer 9/10) at a concentration of approximately 5 mg protein/mL and solubilized by the addition of final concentration 1% 3-(laurylamido)-*N,N*’-dimethylaminopropylamine oxide (LAPAO) ^42^. The sample was inverted several times and was kept on ice for 30 minutes. In the following centrifugation for 30 minutes at 310,000 × g, the cell debris was eliminated. The supernatant (containing the solubilized protein) was incubated with Ni-NTA resin which was pre-equilibrated with 10% glycerol, 600 mM NaCl, 20 mM Tris-HCl (pH 8.0), 5 mM β-mercaptoethanol, 10 mM imidazole, 0.2% LAPAO, and 10 mM melibiose (buffer A) at a concentration of about 25 mL resin/g membrane vesicle protein. After incubation for 1 hour under gentle agitation, the resin with the adsorbed material was centrifuged at 4000 × g (Beckman CoulterTm, USA), washed once with the same medium, and loaded onto with a volume of 10 mL column (Bio-Rad, Lab., Inc.) and buffer exchange done manually. Add 10 mL of buffer A to wash column again. 10 mL buffer B (10% glycerol, 600 mM NaCl, 20 mM Tris-HCl (pH 8.0), 5 mM β-mercaptoethanol, 10 mM imidazole, 0.1% DDM, and 10 mM melibiose) was added to detergent exchange. Repeat 5 times more, don’t mix volume of each washing. Total volume buffer B is 60 mL. And then 10 mL buffer C (10% glycerol, 100 mM NaCl, 20 mM Tris-HCl (pH 8.0), 5 mM β-mercaptoethanol, 10 mM imidazole, 0.1% DDM, and 10 mM melibiose) was added to column to decrease the concentration of NaCl. Repeat 2 times. A final wash of 30 mL was applied before rising the imidazole concentration to 100 mM (eluted buffer) to allow adsorbed proteins to detach from the column.

After evaluation of the flow-through by means of a D_2_-lamp at 280 nm (Varian Cary UV/Vis Spectrophotometer), the fractions containing proteins were pooled. The concentration of the new purification protein was measured using flow formula:

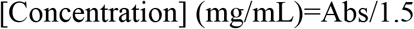

All protein aliquots were put together in flacon flask, using liquid N_2_, and stored at −80°C until further use.

### Reconstitution

Cys-less and R149C mutant permease (≈0.5 mg/mL) solubilized in DDM (0.1%, w/w) were mixed with *E.coli* lipids to give a protein-to-lipid ratio of 1:2 (w/w) and incubated for 10 minutes at 0-4°C. Bio-beads SM-2, which were pre-washed with distilled water and pre-equilibrated with 20 mM MES (pH 6.6) containing 100 mM KCl were stepwise added at a concentration of 120 mg/mL solution to absorb the detergent and the sample was incubated overnight at 4°C ^43^. The proteoliposomes were washed twice in a 100 mM KCl, 20 mM MES buffer (pH 6.6). Proteoliposomes were collected by centrifugation at 310,000 × g for 30 minutes (SORVALL^®^ Centrifuges, USA). Finally, proteoliposomes were resuspended in 100 mM KCl, 20mM MES buffer (pH 6.6), freezed using liquid N2, and stored at −80°C.

### Determination of protein purity

Polyacrylamide gel electrophoresis (12 %) was used to determine the purification of the protein flow through Ni-column and the reconstitution proteoliposome. (Fig.2). Proteoliposome quantity was assayed according to Lowry ^44^ in the presence of 0.2% SDS using bovine serum albumin (BSA) as standard.

**Figure 2.**
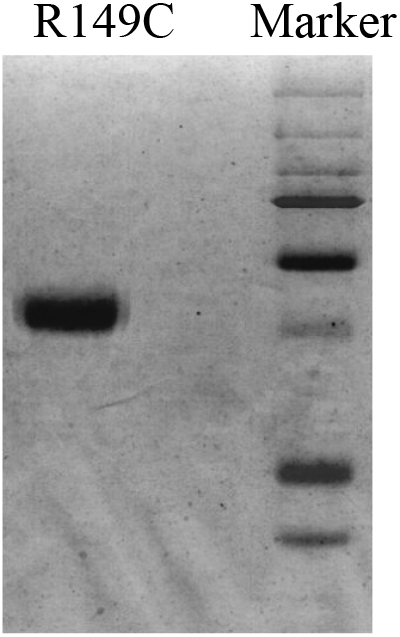
SDS-PAGE of R149C mutant after reconstitution.

**Figure 3.**
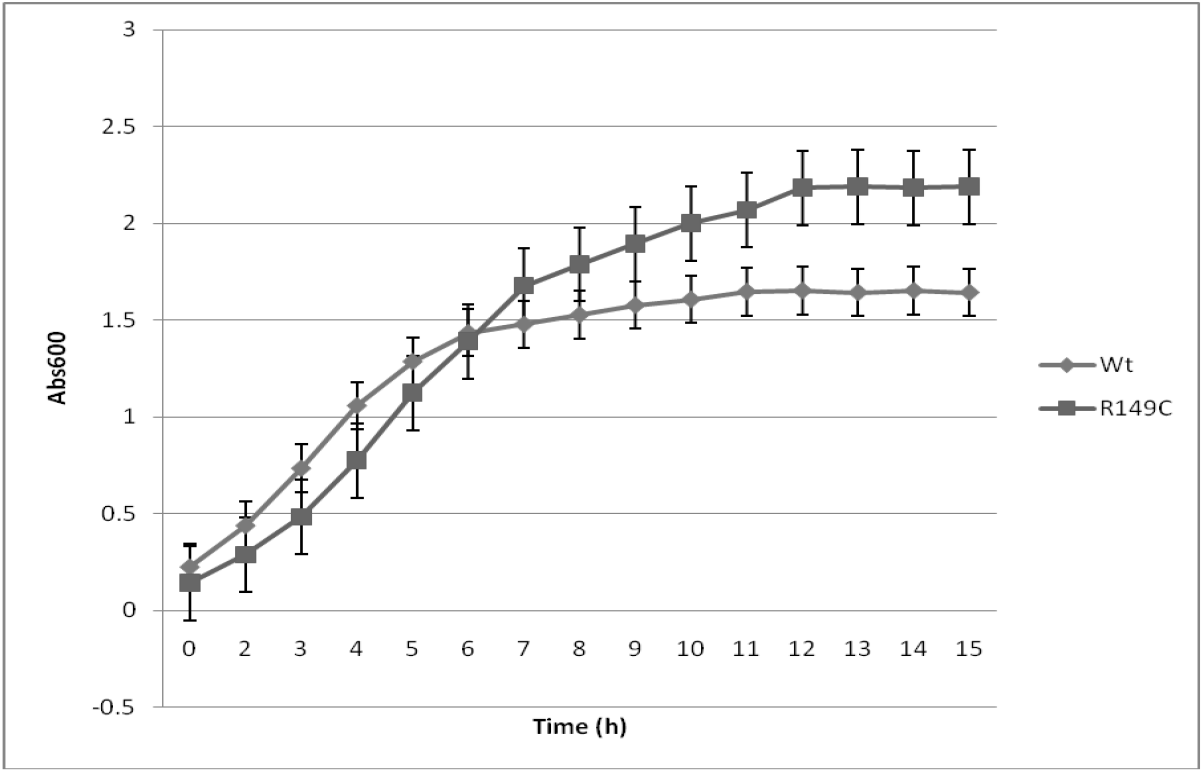
Growth kinetics of R149C and Cys-less cells.

**Figure 4.**
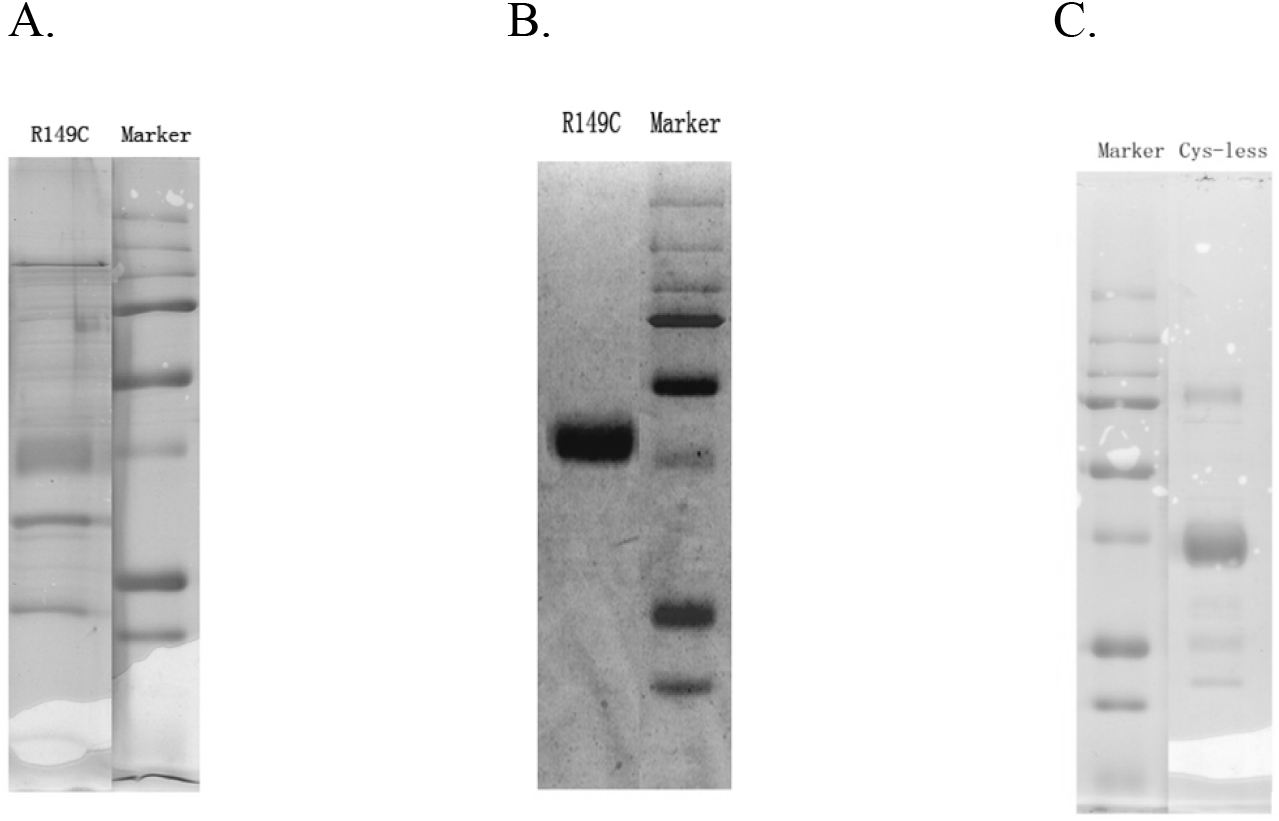
SDS-PAGE of R149C and Cys-less reconstituted protein. A. R149C reconstituted protein in the absence of protease inhibitor; B. R149C reconstituted protein in the present of protease inhibitor; C. Cys-less reconstituted protein.

### Cells growth kinetics determination

Cys-less and R149C cells were picked, put in 12 mL LB medium, and incubated overnight at 30 °C. 3 mL these Mini-culture were transferred in 50 mL M9 medium, and incubated in the Thermoshaker at 30 °C. OD_600nm_ was measured every one hour.

### Sample preparation and FTIR spectra acquisition

Experimental setup was the same as that described in León et al. 2005 with some modifications. A sample of 20 μL of a proteoliposome suspension (about 150 μg of protein) was spread homogeneously on a germanium ATR crystal (50 ×10 × 2 mm, Harrick, Ossining, NY, yielding 12 internal reflections at the sample side) and dried under a stream of nitrogen. The substrate-containing buffer and the reference buffer (containing or not MelB substrates) were alternatively perfused over the proteoliposome film at a rate of ≈1.5 mL/min. The film was exposed to the substrate containing buffer for 2 minutes (4 minutes when the substrate-containing buffer had 10 mM melibiose) and washed with the reference buffer for 10 min (15 min when the substrate-containing buffer had 10 mM melibiose). The switch of buffers was carried out by a computer-controlled electro valve. A Bio-Rad FTS6000 infrared spectroscopy instrument (Hercules, CA) was used for spectra acquisition. For difference spectra induced by Na^+^, each cycle 500 scans at a resolution of 4 cm^−1^ were recorded and a total of 60 spectra were taken and averaged in order to increase the signal-to-noise ratio, i.e., a total of 30,000 scans for every difference spectrum. For difference spectra induced by melibiose, each cycle 1000 scans at a resolution of 4 cm-1 were recorded and a total of 40 spectra were taken to average. Each experiment took about 17 h to be completed, and three separate experiments using newly prepared films were done for each condition. Sample temperature was adjusted to 20°C using a cover jack placed over the ATR crystal and connected to a circulating thermostatic bath. The cover jack temperature was controlled with a fitted external probe.

### Data corrections

Spectra corrections were essentially carried out as described previously ^28^. water difference spectrum induced by the substrate(s), absorbance of the substrate(s) (in our case melibiose, since the cations do not absorb), and change in the swelling of the film, with an apparent gain/loss of water with a concomitant apparent loss/gain of sample (protein and lipid) may be contribute to the experimental difference spectrum. The latter contribution was corrected by subtracting, from the experimental difference spectrum, an absorption spectrum of MelB proteoliposomes in the substrate-containing buffer. The subtraction factor used was that able to remove the lipid CH_2_ bands at 2920 and 2850 cm^−1^. Water difference spectrum induced by substrate and absorbance of substrate were corrected subtracting from the experimental difference spectrum a difference between the substrate-containing buffer and the reference buffer. The subtraction factor was adjusted to flatten the water band between 3700 and 2800 cm^−1^ and to remove bands coming from the substrates (melibiose give an intense band at 1080 cm^−1^, whereas cations give no bands).

### Spectra deconvolution

Deconvolution by the maximum entropy method was applied to difference spectra to resolve overlapped bands. Deconvolution strongly enhances the noise in the data. Therefore, for being applicable to real data, deconvolution must operate with some noise suppression. The most habitually used deconvolution method is Fourier deconvolution (FD), which discriminates between signal (noiseless data) and noise only on the basis of their frequency. Recently, Lorenz-Fonfria and Padros 2005 have introduced a new entropy expression that is particularly suited for the deconvolution of difference spectra. Deconvolution was performed using a Lorentzian band of 7 cm^−1^ width. This value was determined from the spectra as described. For the maximum entropy deconvolution, the same regularization parameter (10^−11^) was used for all spectra. Its value was derived in accordance with the spectral noise content.

### Intrinsic fluorescence measurements

Fluorescence measurements were performed using QuantaMaster™ UV VIS spectrofluorometer, and were processed with Felix 32 software (Photon Technology International, Lawrenceville, NJ, USA). All the experiments were carried out at 20°C. Measurements were carried out on samples containing 20 μg of protein/mL that were previously dissolved at 100 mM KPi pH 7.0, sonicated for 30 seconds (Ultrasonic cleaner of Fungilab s.u., Spain) and placed in 1.5 mL cuvette (Quartz glass, 10 mm light path). A special component on bottom of the cuvette holder allows efficient mixing with small magnets. Each experiment was performed in triplicate. Data were corrected by subtracting the spectra of buffer. All the buffers were cleared using 0.45 μm filter.

## Results

### Cells growth, protein expression and purification

The vector pK95ΔAHB, which express R149C or Cys-less mutant permeases was transfected into *E.coli* DW2-R and verified by sequencing (data not show). After overnight growth, the OD_600nm_ of cells which express R149C mutant permease (R149C cells) reach about 1.8-2.2, whereas OD_600nm_ of cells which express Cys-less mutant permease (Cys-less cells) will be 1.0-1.5. More R149C cells were harvested under same condition. In the first instance, we analysed the ability of R149C and Cys-less cells to grow over a period of 0-15 h. 2μl cells (stored at −80°C) were plated on MacConkey agar plates (see Materials and Method) and incubated overnight at 30°C. Red small colonies were found for Cys-less cells, whereas the colonies of R149C cells were white. Fig.3 shows the growth kinetics of Cys-less and R149C cell. During the first six hours, OD600 of R149C is lower than Cys-less. However, six hours later, the OD600 of R149C is higher than Cys-less.

In the course of purification, the protease inhibitor AEBSF was added to the buffer to avoid the effect of proteases. Fig.4 shows the effect of AEBSF on the purification of R149C. Replacement of Arg-149 by neutral Cys residue may led to a protease-sensitive MelB. If no protease inhibitor was added during purification, only a very tiny protein was obtained (Fig.4 A). Most protein was degraded by proteases, as indicated by the bands seen under R149C band. Compared to R149C mutant permease, Cys-less seem more stable (Fig.4 C). Under the same conditions, 5 mg of the Cys-less protein were obtained from 16 L culture, whereas only 0.6 mg of the R149C protein were obtained (Fig. 5). Later, we measured the infrared spectra of these purified protein, which were reconstituted into the *E.coli* lipids. The infrared spectra of R149C was very different to Cys-less (data no show). Later on, R149C was purified and reconstituted in the present of protease inhibitor (Fig.4 B). In this case, more protein (about 4 mg) was obtained from 16 L culture, which was about 80% of Cys-less under same conditions. From Fig.4 B we can see that there are no degradation bands under R149C band. In this case, the infrared absorption spectrum was similar to that of Cys-less.

**Figure 5.**
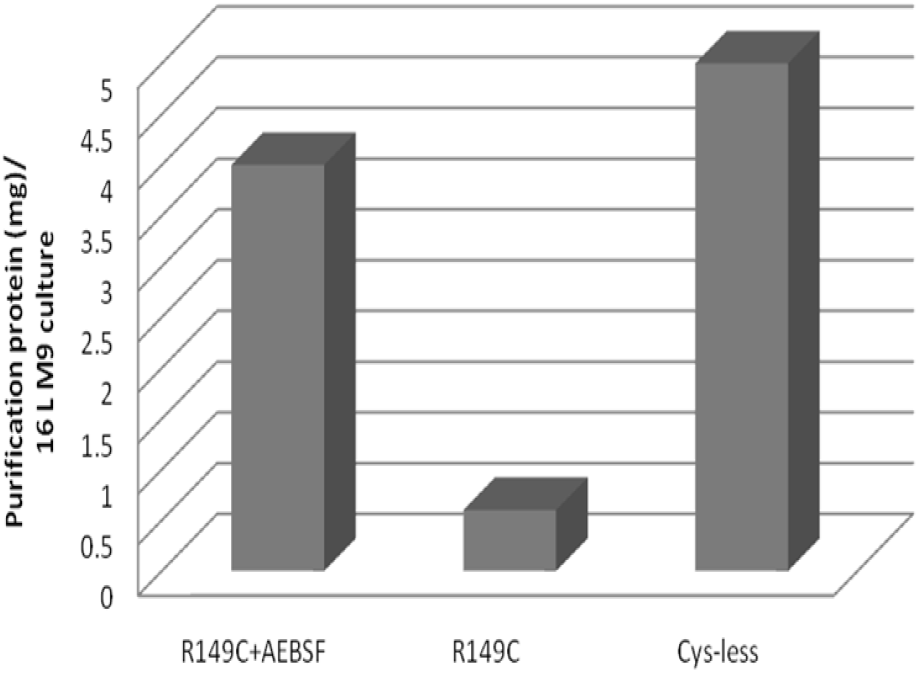
Comparison of the yield of purified protein of R149C and Cys-less mutant permeases.

### Substrate-induced changes of the Cys-less mutant

MelB mutant was constructed using Cys-less as molecular background. León et al. has checked the similarity of the substrate-induced difference spectra of the Cys-less and the WT permeases (León et al. 2009). Peaks absorbing in turns, β-structures, α-helices, and carboxylic side-chain regions are observed in the difference spectra of both permeases. All respective difference spectra are similar in shape and share the same main peaks. It can be concluded that the substrate-induced difference spectra of Cys-less and WT are essentially similar, albeit not entirely identical, and yield comparable information on fully active MelB permeases.

Since the difference infrared spectra of Cys-less and WT permeases are similar, as are their behavior, Cys-less mutant permease was selected as a control in this study. In all experiments, the Na^+^ and/or sugar concentrations used (10 mM) are saturating (Bassilana et al. 1988; Damiano-Forano et al. 1986). Data acquisition and spectra deconvolution have been presented at Materials and Methods. Comparing our data with former published spectra, the main parts of the difference spectra of Cys-less and WT are very similar, i.e., the amide I region in Na^+^ versus H^+^ and melibiose·Na^+^ versus Na^+^ difference spectra, the carboxylic peaks around 1400 cm^−1^ in Na^+^ versus H^+^ difference spectrum, etc (León et al. 2006 and León et al. 2005).

### Substrate-induced changes of the r149c mutant

The difference spectra induced by Na^+^ binding to R149C and Cys-less are similar, however, there are many alterations in the amide I and II region (Fig. 6A). This means that the structural changes resulting from the replacement of the coupling H^+^ by Na^+^ are a little alike in both permeases. Some variations are visible in some parts of the difference spectrum, such as those peaks in the 1630–1600 cm^−1^ region. These changes may be mainly caused by the intensity of the peak centered at 1613 cm^−1^, which may be assigned to Arg side chains and more specifically to Arg149. The assignment of this peak to Arg side chains is consistent with the fact that this peak is sensitive to H/D exchange. In addition, there is a decrease in the intensity of the peaks absorbing in the amide II (around 1550 cm^−1^). Peaks corresponding to carboxylic acids (at 1599–1600, 1571–1576, 1404, and 1383 cm^−1^) show no modifications except for a disappearance in the peak at (-)1383 cm^−1^. There are some variations in the peaks in the 1650-1700 cm^−1^ region. These changes may be mainly caused by the disappearance of the peak centered at 1672 and 1657 cm^−1^. The peak at 1672cm^−1^ may be assigned to Arg side chains (Baenziger et al. 1997). Some noteworthy dissimilarities are at 1663 and 1653 cm^−1^. The peak at 1663 cm^−1^ may correspond to signal variation arising either from α-helix or from loops. Peaks detected at 1658 and 1651 cm^−1^ may be attributed to 2 distinct types of helical structures (type I and type II, respectively). A comparison of the deconvoluted spectra (Fig.6 B) that increase the peak resolution confirms these conclusions.

**Figure 6.**
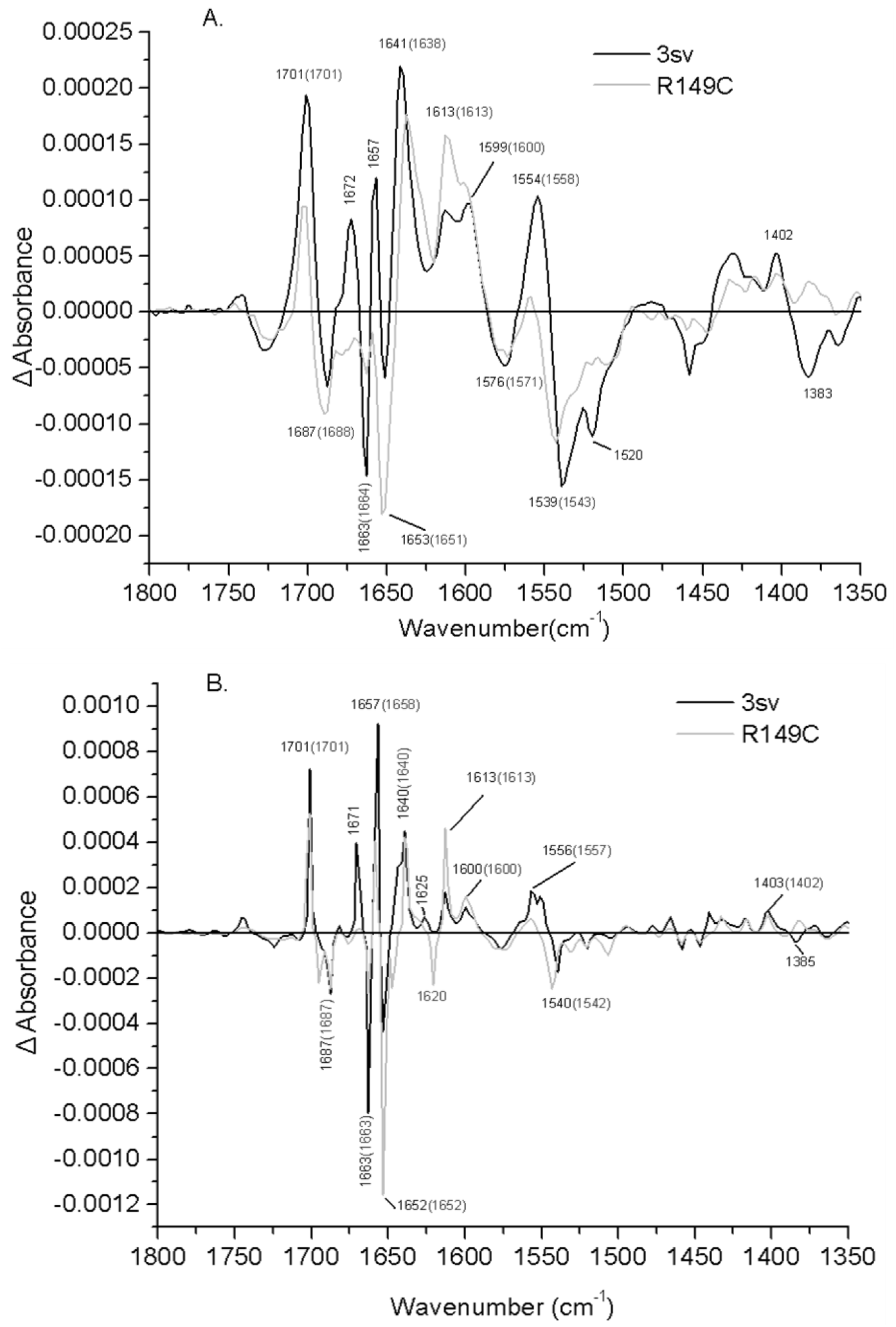
Comparison of Na^+^-induced difference spectra of R149C and Cys-less. (A) gray line: difference spectrum of R149C in 20 mM MES, 100 mM KCl, 10 mM NaCl, pH 6.6, minus R149C in 20 mM MES, 110 mM KCl, pH 6.6; black line: difference spectrum of MelB Cys-less under the same conditions. (B) Deconvoluted difference spectra of panel A.

The R149C difference spectrum induced by melibiose binding in the presence of Na^+^ (Fig. 7 A) presents an overall shape comparable to that of Cys-less. However, there are some noteworthy dissimilarities in the amide I region. There is a decrease in the intensity of the peaks absorbing around 1668 cm^−1^, not seen in the difference spectrum due to Na^+^ binding, is expected to reflect changes in turns or in open loop signals. Another dissimilarity is a decrease in the intensity of the peaks 1643 cm^−1^, which is best seen after deconvolution of the spectra (Fig.7 B). This peak centered at 1643 cm^−1^ may correspond to the changes in β-sheets, 3_10_ helices or open loops. In addition, there is an increased intensity of the peaks absorbing at 1651 cm^−1^, which suggest changes in α-helix signals. Therefore, there are some changes in α-helix when R149C binds melibiose compared to Cys-less.

**Figure 7.**
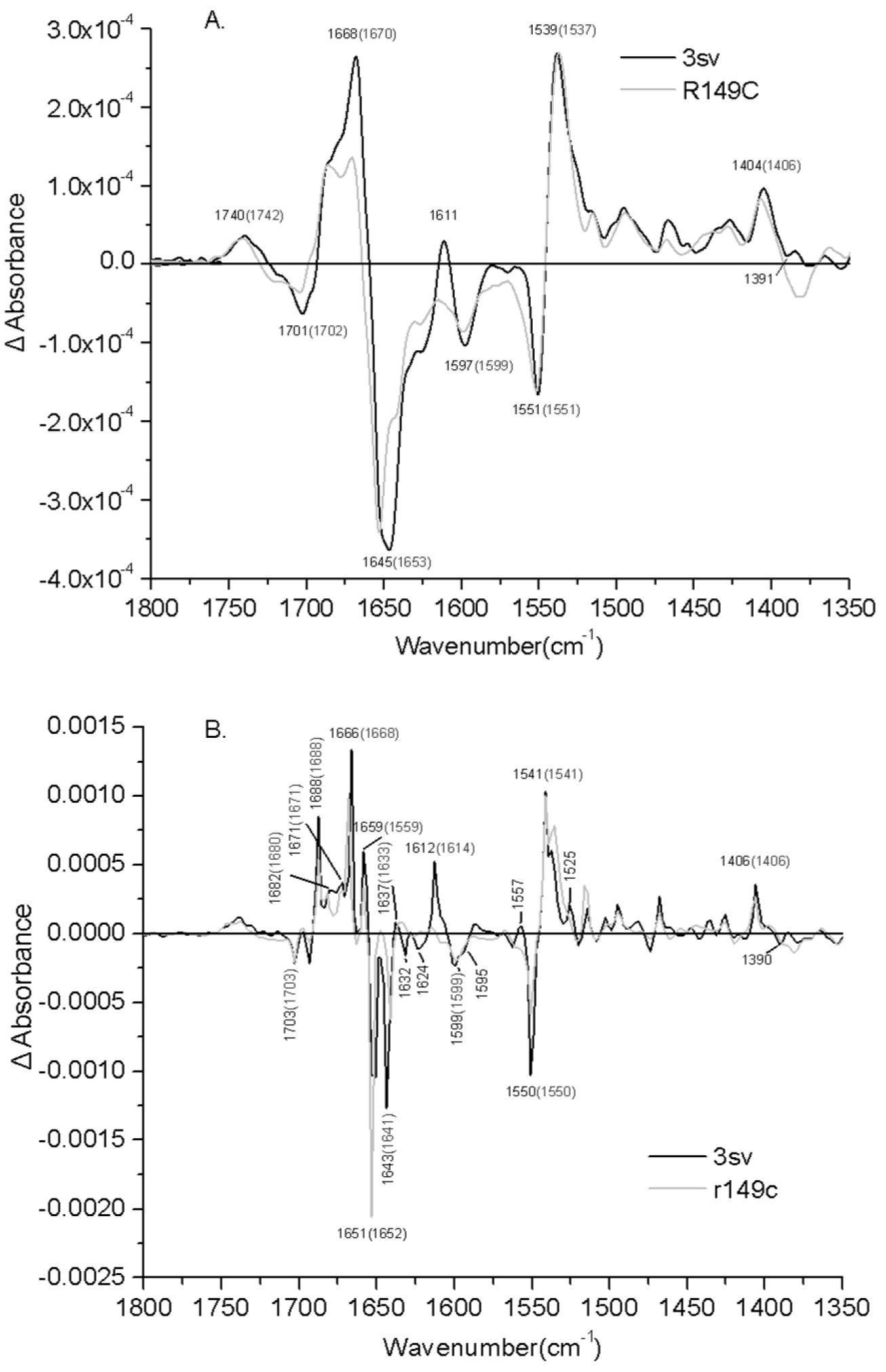
Comparison of melibiose-induced difference spectra of R149C and Cys-less in the presence of Na^+^. (A) gray line: Difference spectrum of R149C in 20 mM MES, 100 mM KCl, 10 mM NaCl, 10 mM melibiose, pH 6.6, minus R149C in 20 mM MES, 100 mM KCl, 10 mM NaCl, pH 6.6; black line: difference spectrum of MelB Cys-less under the same conditions. (B) Deconvoluted difference spectra of panel A.

The peaks at 1688, 1682, and 1671 cm^−1^ of Cys-less, previously assigned to putative β-structures or turns, are also found at similar position in R149C. This suggests that the conformational changes of these structures occurring in R149C upon melibiose interaction are similar from those of Cys-less. The 1659 cm^−1^ peak, which was previously proposed to arise from one of the two major α-helix subpopulations (α-helix type I) also is found in R149C mutant. However, it could not be found in the R141C mutant. It seems that R149C and Cys-less produce some similar secondary structure changes when binding melibiose. There is a decrease at 1693 cm^−1^, which may correspond to changes in β-sheets or side chains. Other changes seen in the deconvoluted amide I region include the disappearance of the negative peak at 1632 cm^−1^ (Fig. 7 B). This spectral variation is consistent with changes in the contribution of β-sheets involved in cooperative interaction between the two substrate-binding sites or in the substrate translocation process. A decrease in the intensity of the peaks near 1624 cm^−1^ in the difference spectra induced by melibiose binding in the presence of Na^+^ or H^+^ (Fig. 9 B) could be interpreted as the disappearance of the signal arising from one or more of the side chains known to be involved in the transport function (Arg52, Arg141, and Arg149).

In the amide II region, only small differences are observed between both permeases, and they can be assigned mainly to amino acid side chains. There is the absence of the 1525 cm^−1^, which is ascribed to Lys side chains, although it could also correspond to amide II vibration. In the R149C difference spectrum (Fig. 7 B) the peak centered at 1541 cm^−1^ (assigned to amide II) is very similar to the peak of Cys-less. Finally, carboxylic peaks around 1703, 1599, and 1406 cm^−1^ are comparable in the difference spectrum (Fig. 7 B).

In summary, the observed variations of sugar-induced FTIR signals at the level of α-helices, as well as from turns and β-sheets between the R149C and Cys-less permeases, indicate that the Arg149 mutation has an impact on the MelB structural variations associated with or triggered by sugar binding.

### Intrinsic tryptophan fluorescence spectrum of MelB Cys-less mutant

The intrinsic fluorescence properties of MelB Cys-less and R149C mutant permeases were first compared in the absence of ligands (Figure 8). Figure 8 shows the emission fluorescence spectrum recorded upon illumination (290±5 nm) of a proteoliposome suspension prepared from the Mel-6His (Cys-less or R149C mutant) permease-rich fraction (90% permease) and also illustrates the typical spectral changes elicited by successive addition of NaCl and melibiose. In the absence of substrates, the proteoliposome fluorescence spectrum (spectrum 1) is characterized by a maximum emission wavelength centered around 325 nm. NaCl was then added at a final concentration (10 mM) known to produce maximal activation of the sugar-binding and transport activities of the permease. This addition induces general and limited quenching of the fluorescence spectrum for Cys-less (≈2%, spectrum 2, Fig. 8 a) and R149C (≈3.9%, spectrum 2, Fig. 8 b) with no concomitant shift of the maximum emission wavelength. Melibiose was then added at a concentration (10 mM) at which binding and transport activities are maximal. This addition produces a broad increase in fluorescence emission for Cys-less (≈20%, spectrum 3, Fig.8 a) and R149C (≈11%, spectrum 3, Fig.8 b).

**Figure 8.**
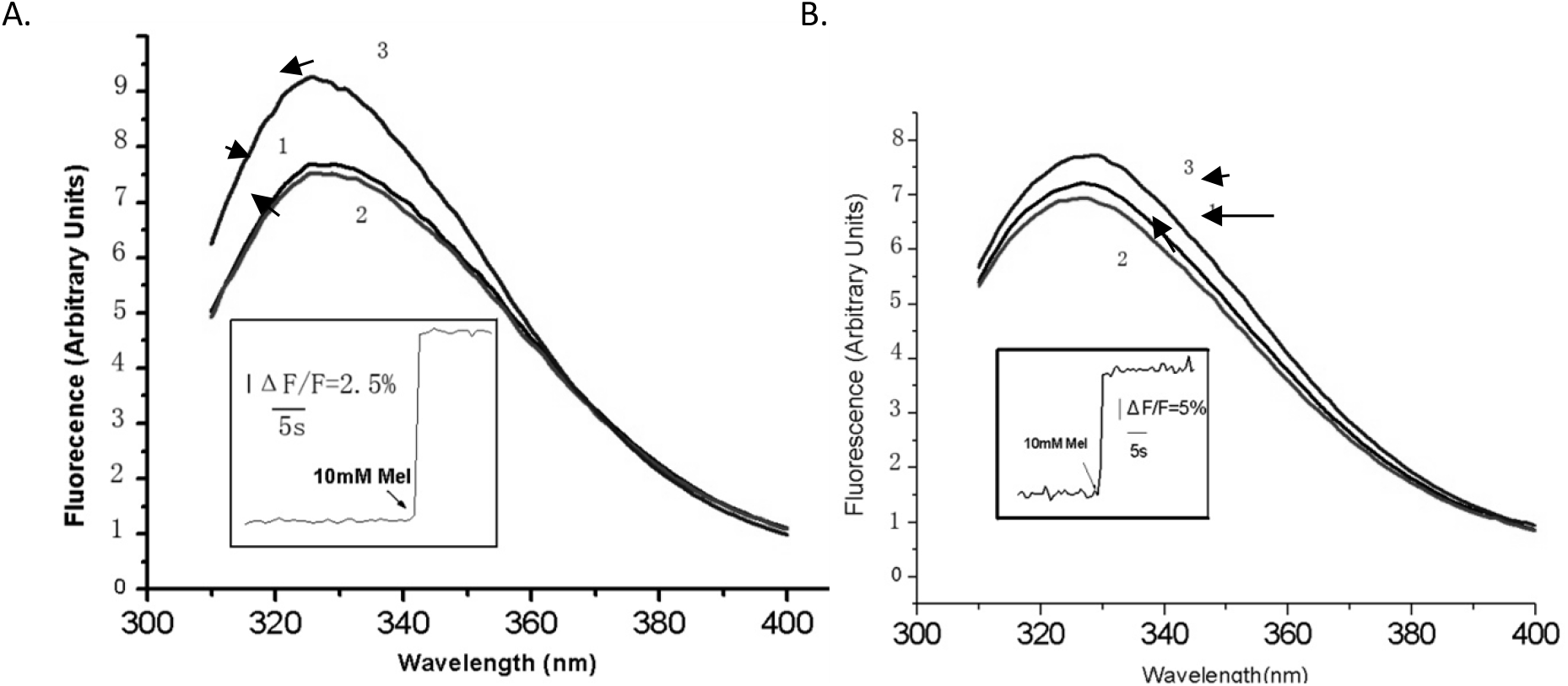
Effects of sodium and melibiose on the intrinsic fluorescence of proteoliposomes containing MelB-6His permease. (A) Cys-less mutant; (B) R149C mutant. A permease-rich fraction containing 90% MelB-6His (Cys-less or R149C mutant) permease was reconstituted into liposomes using total *E.coli* lipids as described in Material and Methods. Samples (2 ml, 20 μg of protein/ml) equilibrated in 0.1 M KPi (pH 7.0) were illuminated at 290±5nm, and the fluorescence emission spectrum was recorded between 310-400 nm. The maximum emission wavelength is at around 325 nm (spectrum 1). NaCl and melibiose were successively added to a final concentration of 10 mM, and the effects on the fluorescence spectra were recorded (spectra 2 and 3, respectively). All spectra were corrected for inner filter effects using the signal measured from a pure liposome preparation with identical absorbancy. Inset: Effect of the addition of sugar on the integrated fluorescence signal recorded from proteoliposome samples (20 μg of protein) equilibrated in 0.1 M KPi (pH 7) and 10 mM NaCl. After excitation at 290±5 nm, the emitted fluorescence light (*F*) was integrated between 310 and 400 nm and recorded as a function of time. The sugar-induced change (Δ*F*) in signal was expressed as Δ*F/F* (in %). Arrow: addition of the indicated sugar at a final concentration of 10 mM. All experiments were carried out at 20°C.

**Figure 9.**
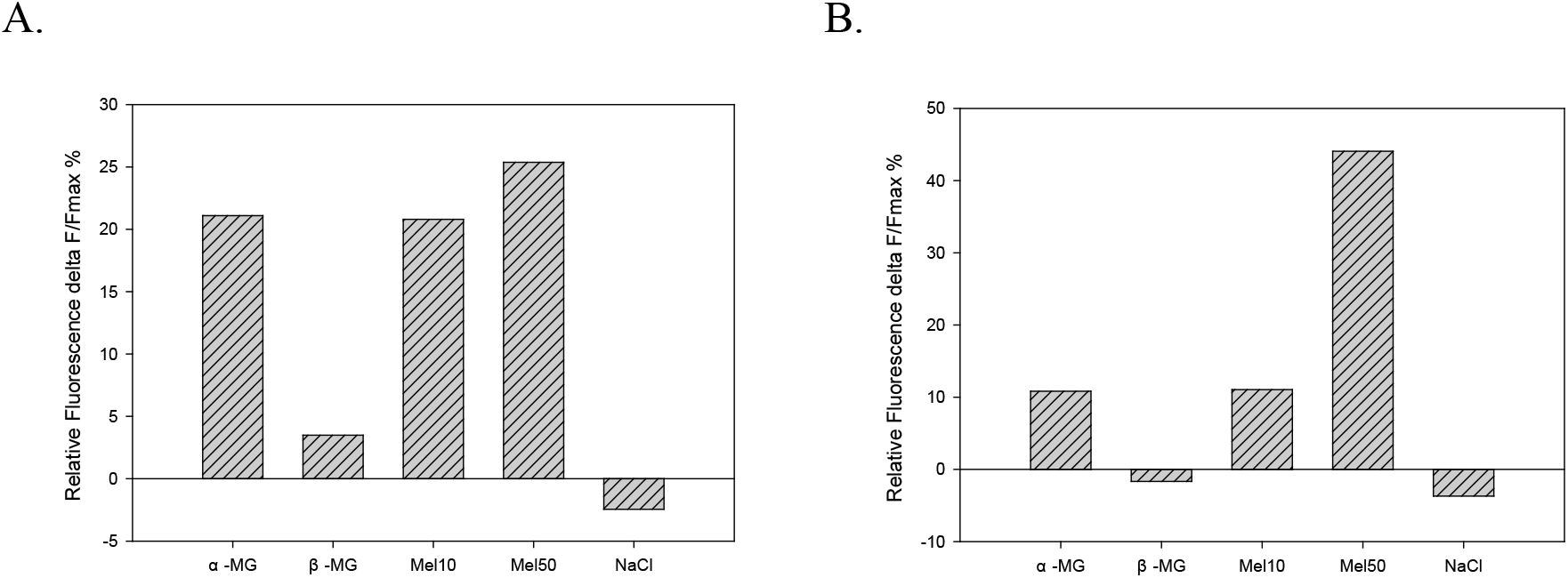
Effect of different galactosides on the tryptophan fluorescence of MelB Cys-less (a) and R149C (b). Proteoliposomes were equilibrated in 100 mM KPi, 10 mM Tris-HCl (pH 7.0), and 10 mM NaCl. Samples (1.5 ml, 20 μg of protein/ml) were incubated at 20°C and illuminated at 290 nm, and the relative variation in fluorescent light emitted between 310 and 400 nm was recorded. Each sugar was added at a final concentration known to induce maximal variation in the fluorescence signal. The following additions were made: methyl α-D-galactopyranoside (α-MG, 15 mM), methyl β-D-galactopyranoside (β-MG, 35 mM), melibiose (Mel10, 10 mM), and melibiose (Mel50, 50 mM), NaCl (10 mM).

Interpretation of the changes in the fluorescence spectrum of Mel-6His permease induced by the substrates is complex. The carrier intrinsic fluorescence properties were therefore analyzed by measuring the relative variation in the fluorescence signal (Δ*F/F*) integrated between 310 and 400 nm. Figure 8 (inset) shows that, in the presence of a saturating concentration of sodium (10 mM), the fluorescence signal increases by 20% for Cys-less and 11% for R149C upon addition of the physiological substrate melibiose.

### Effect of sugars on the mutant fluorescence properties

Mus-Veteau and coworkers reported previously that addition of 10 mM NaCl to proteoliposomes containing WT permease quenched the intrinsic fluorescence by about 2%. Similar limited signal quenching is observed with Cys-less and R149C mutant permeases constructed in this study (for Cys-less 1.8%, for R149C 3.9%, Fig. 9), indicating that similar conformation change between Cys-less and R149C mutant permease when binding Na^+^.

MelB permease recognizes a broad range of α-and β-galactoside derivatives and, to a lesser extent, galactose. To compare the effects of various sugar substrates on MelB permease Cys-less and R149C mutant fluorescence properties, proteoliposomes were equilibrated in the presence of a saturating concentration of sodium (10 mM), and the concentration of each sugar was added at a concentration giving a maximal fluorescence change (ΔF/Fmax). Figure 9 shows that saturating concentrations (10-50 mM) of α-galactosides such as α-MG, melibiose produced a large ΔF/Fmax. The melibiose (10 mM)-dependent ΔF/Fmax change is about 20.8% of Cys-less. However, melibiose (10 mM) induces smaller change of R149C mutant compared to Cys-less at the same condition. The melibiose (10 mM)-dependent ΔF/Fmax change is about 11.0% (Fig.9 b). 50mM melibiose was used to induce more fluorescence change of Cys-less and R149C mutant. For Cys-less, when more melibiose was added, only a small increase in the fluorescence was found. The melibiose (50 mM)-dependent ΔF/Fmax change is about 25.7% (Fig.9 a). However, when the concentration of melibiose was change from 10 mM to 50 mM, the melibiose-dependent ΔF/Fmax change increases from 11.0% to 44.1% (Fig.9 b). The maximal ΔF/F produced by β-galactosides such as the monosaccharide β-MG was at best only 7% and necessitated a higher sugar concentration (>35mM) (Mus-Veteau et al., 1995). There is a similar result for Cys-less and R149C mutant permeases. It was finally shown that β-MG produces a ΔF/Fmax of 3.5% of Cys-less, whereas the maximal ΔF/F produced by β-MG of R149C is −1.7%. So this β-galactoside has no effect of R149C because this intensity variation corresponds to the dilution effect.

## Discussion

Former studies suggest that mutation of Arg149 into Cys or Asn (but not Lys) leads to inhibition of α-NPG (*p*-nitrophenyl α-D-galactopyranoside) binding. Dayem et al. had provided evidence that mutagenesis of two basic residues, Arg141 and Arg149, located in the inner loop 4-5 of MelB permease from *E.coli* impaired the Na^+^-sugar symporter function. The data suggest that Arg-141 primarily takes part in the reaction of co-substrate translocation, whereas the nearby Arg149 residue may be located close or at the sugar binding site and most likely contributes to its structural organization. Due to the absence of a high-resolution 3D structure of MelB, the presence of loops or the localization of amino acids is still tentative and is based on different kinds of studies. Recently obtained 3D structures of some transporters such as the lactose permease (lacY, 12 transmembrane helices) may help in recognizing the structure and function of MelB. Yousef and Guan reported a 3D structure model of the melibiose permease of Escherichia coli threaded through a crystal structure of the lactose permease of *E.coli* (LacY), manually adjusted, and energetically minimized. The 3D model indicates that MelB consists of two pseudosymmetrical 6-helix bundles lining an internal hydrophilic cavity, which faces the cytoplasmic side of the membrane. Both sugar and cation binding sites are proposed to lie within the internal cavity. The 3D model also suggests that the salt bridge between Arg149 (helixV) and Asp 124 (helix IV) may be is essential for sugar binding and selectivity. In this study, we first applied attenuated total reflection Fourier transform infrared (ATR-FTIR) difference spectroscopy to obtain information about the structural changes involved in substrate interaction with the R149C mutant and with the MelB Cys-less mutant. Our data indicate that only limited modifications of the substrate-induced difference spectral features take place in R149C compared with Cys-less. The conclusion emerges that small structural rearrangements occurring in key sites in the permease are sufficient to produce such significant functional effects.

The R149C mutant cannot transport substrates, but our data indicate that R149C mutant retains the ability to bind them (see intrinsic fluorescence evidences, shown in the Result part, and discussion below). The Na^+^-induced difference spectra for R149C and Cys-less indicate that replacement of H^+^ by Na^+^ as a coupling ion induces very similar conformational changes in the amide I region and the carboxylic side-chain absorption region. This means that substrates induce similar conformational changes when they bind to the R149C and Cys-less. However, some noteworthy dissimilarities are at 1663, 1658, and 1651 cm^−1^. The peak at 1663 cm^−1^ may correspond to signal variation arising either from α-helix or from loops. Peaks detected at 1658 and 1651 cm^−1^ may be attributed to 2 distinct types of helical structures (type I and type II, respectively) (Fig.6). All these different changes suggest that R149C mutant modifies the binding features of Na^+^ ion. The permease intrinsic fluorescence variations also show the same conclusion. When 10 mM NaCl was added to proteoliposomes containing Cys-less permease, a quenching of 1.8% of the signal is observed (Fig. 8A). However, a quenching of 3.9% is observed with R149C mutant permease in the same conditions (Fig. 8B). So Arg-149 is also very important to configure properly the melibiose permease cation binding site.

The melibiose-induced difference spectra for R149C and Cys-less is more similar in the amide II and the carboxylic side-chain absorption region (Fig. 9 A). However, there are some noteworthy dissimilarities in the amide I region. The decrease in the intensity of the peaks absorbing around 1669 cm^−1^, not seen in the difference spectrum due to Na^+^ binding, is expected to reflect changes in turns or in open loop signals. Another big dissimilarity is a decrease in the intensity of the peak at 1643 cm^−1^, which is best seen after deconvolution of the spectra (Fig.9 B). This peak centered at 1643 cm^−^ 1 may correspond to the changes in β-sheets, 3_10_ helices or open loops (Dave et al. 2000). In addition, there is an increased intensity of the peaks absorbing at 1653 cm^−1^, which suggest changes in α-helix signals. These data indicate that secondary structures suffer different changes in R149C and Cys-less upon melibiose binding. All of this structural information is in agreement with the idea that Arg-149 residue plays a very important role in sugar binding or close to the sugar binding site.

Melibiose permease recongnizes a broad range of α- and β-galactoside derivatives and, to a lesser extent, galactose. To compare the effects of various sugar substrates on MelB R149C and Cys-less fluorescence properties, different sugars were incubated with proteoliposomes, and the concentration of each sugar giving a maximal fluorescence change (ΔF/Fmax) was determined. Saturating concentrations of α-galactosides such as the monosaccharide αMG, or the disaccharide melibiose produced a large ΔF/Fmax, whereas the ΔF/Fmax produced by β-galactosides such as the monosaccharide βMG was at best only 7% for Cys-less, however, was −1.67% for R149C. All these data indicate that the intrinsic fluorescence variations properties of R149C and Cys-less are very similar. However, R149C mutant seem more sensitive to the sugar. Mus-Veteau et al. had indicated a correlation between the amplitude of fluorescence changes produced by a given sugar and the permease affinity for this sugar. Replacement of Arg-149 by neutral residues (Cys) affect the sugar selectivity and binding, showing higher affinity to α-galactosides, however, lower affinity to β-galactosides. From these data, we speculate Arg-149 may play an important role in sugar selectivity and binding. Replacement of Arg-149 by neutral residues (Cys) will change the salt bridges between Arg-149 (helix V) and nearby residues (for example Asp-124 in helix IV), which are strictly conserved among all MelB orthologues, then changing the structure of MelB internal hydrophilic cavity.

## Conclusions

Replacement of Arg-149 by Cys produces a permease that is still able to bind Na^+^ and melibiose as indicated by the difference infrared spectrum and tryptophan fluorescence variations. The conformational changes of R149C and Cys-less permeases induced by Na^+^ and melibiose binding are similar, however with several dissimilarities especially for melibiose binding. Arg149 is probably involved in the structuring of the substrates binding site and probably participates also in the release mechanism of substrates, since there is no transport but there is substrate binding.

## Ethics

Not applicable

## Authors’ contributions

YL designed the experiments, carried out the practical work and drafted the manuscript.

